# FUMA: Functional mapping and annotation of genetic associations

**DOI:** 10.1101/110023

**Authors:** Kyoko Watanabe, Erdogan Taskesen, Bochoven Arjen van, Bochoven Arjen van, Danielle Posthuma

## Abstract

A main challenge in genome-wide association studies (GWAS) is to prioritize genetic variants and identify potential causal mechanisms of human diseases. Although multiple bioinformatics resources are available for functional annotation and prioritization, a standard, integrative approach is lacking. We developed FUMA: a web-based platform to facilitate functional annotation of GWAS results, prioritization of genes and interactive visualization of annotated results by incorporating information from multiple state-of-the-art biological databases.

---

In the past decade, more than 2,500 genome-wide association studies (GWAS) have identified thousands of genetic loci for hundreds of traits^1^. The past three years have seen an explosive increase in GWAS sample sizes^2–4^, and these are expected to increase even further to 0.5-1 million in the next year and beyond^5^. These well-powered GWAS will not only lead to more reliable results but also to an increase in the number of detected disease-associated genetic loci. To benefit from these results, it is crucial to translate genetic loci into actionable variants that can guide functional genomics experimentation and drug target testing^6^. However, since the majority of GWAS hits are located in non-coding or intergenic regions^7^, direct inference from significantly associated single nucleotide polymorphisms (SNPs) rarely yields functional variants. More commonly, GWAS hits span a genomic region (‘GWAS risk loci’) that is characterized by multiple correlated SNPs, and may cover multiple closely located genes. Some of these genes may be relevant to the disease, while others are not, yet due to the correlated nature of closely located genetic variants, distinguishing relevant from non-relevant genes is often not possible based on association P-values alone. Pinpointing the most likely relevant, causal genes and variants requires integrating available information about regional linkage disequilibrium (LD) patterns and functional consequences of correlated SNPs. Ideally, functional inferences obtained from different repositories are integrated, and annotated SNP effects are interpreted in the broader context of genes and molecular pathways. For example, consider a genomic risk locus with one lead SNP associated with an increased risk for a disease, and several dozen other SNPs in LD with the lead SNP that also show a low association P value, spanning multiple genes. If none of these tested SNPs and none of the other (not tested but known) SNPs in LD with the lead SNP are known to have a functional consequence (i.e. altering expression of a gene, affecting a binding site or violating the protein structure), no causal gene can be indicated. However, if one or several of the SNPs are known to affect the function of one of the genes in the area, but not the other genes, then that single gene has a higher probability of being functionally related to the disease.

In practice, the extraction and interpretation of the relevant biological information from available repositories is not always straightforward, and can be time-consuming as well as error-prone. We have, therefore, developed FUMA, which functionally annotates GWAS findings and prioritizes the most likely causal SNPs and genes using information from 14 biological data repositories and tools (Supplementary Table 1). Results are visualized to facilitate quick insight into the implicated molecular functions. FUMA is available as an online tool at http://fuma.ctglab.nl, where users can set several parameters to filter SNPs or specify specific tissues to be used for annotation based on expression data (Supplementary Table 2 and Supplementary Fig. 1). As input, FUMA takes summary statistics from GWAS.

**Figure 1.**
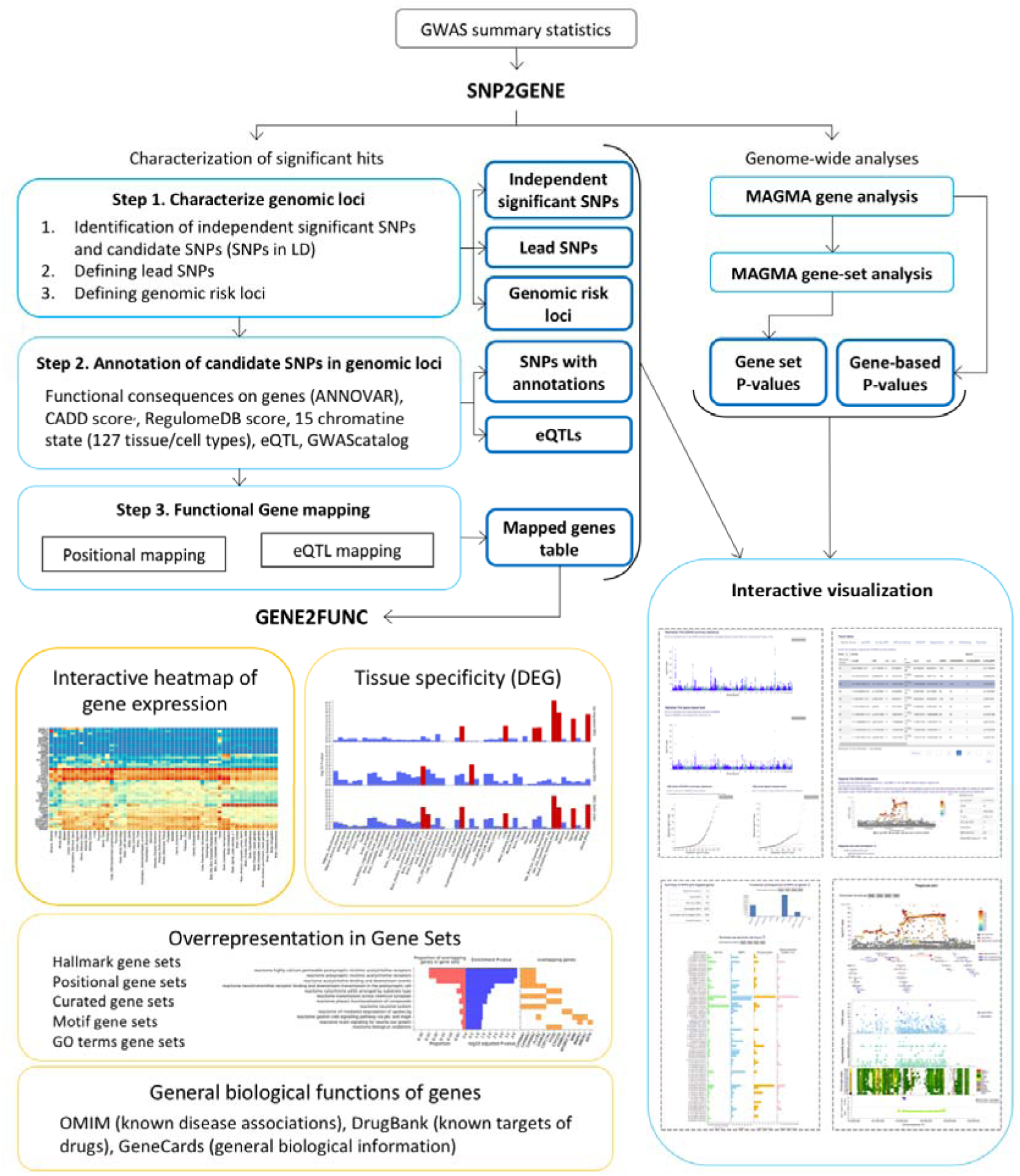
Overview of FUMA. FUMA includes two core processes, *SNP2GENE* and *GENE2FUNC*. The input is GWAS summary statistics. *SNP2GENE* prioritizes functional SNPs and genes, outputs tables (blue boxes), and creates manhattan, quantile-quantile (QQ) and interactive regional plots (box at right bottom). *GENE2FUNC* provides four outputs; a gene expression heatmap, enrichment of differentiallyexpressed gene (DEG) sets in a certain tissue compared to all other tissue types, overrepresentationof gene sets, and links to external biological information of input genes. All results are downloadable as text files or high-resolution images.

The core function of FUMA is the *SNP2GENE* process (Fig.1; Online Methods), in which SNPs are annotated with their biological functionality and mapped to genes based on positional and functional information of SNPs. First, conditional on the provided summary statistics, independent lead SNPs and their surrounding genomic loci are identified depending on LD structure. Lead SNPs and SNPs which are in LD with the lead SNPs are then annotated for functional consequences on gene functions (based on Ensembl genes (build 85) using ANNOVAR^8^), deleteriousness score (CADD score^9^), potential regulatory functions (RegulomeDB score^10^ and 15-core chromatin state predicted by ChromHMM^11^ for 127 tissue/cell types^12,13^) and effects on gene expression using expression quantitative trait loci (eQTLs) of various tissue types (see Online Methods). At this stage, lead SNPs and correlated SNPs are also linked to the GWAS catalog^1^ to provide insight into previously reported associations with a variety of phenotypes. Functionally annotated SNPs are subsequently mapped to genes based on functional consequences on genes annotated by ANNOVAR (positional mapping) and/or eQTLs of user defined tissue types (eQTL mapping). Gene mapping can be controlled by setting several parameters (Supplementary Table 2) that allow to in or exclude specific functional categories of SNPs. For example, positional mapping may optionally use only *coding* SNPs for gene mapping. For eQTL mapping, specific tissues can be selected to only include SNPs that influence the expression of genes in the selected tissue(s) (Online Methods and Supplementary Table 2). By combining positional mapping of deleterious coding SNPs and eQTL mapping across (relevant) tissue types (i.e. functional mapping; Online Methods), FUMA enables to prioritize genes that are highly likely involved in the trait of interest. Due to the use of eQTL information, the prioritized genes – although influenced by SNPs within a disease-associated locus are not necessarily themselves located inside that locus.

To obtain insight into putative causal mechanisms, the *GENE2FUNC* process annotates the prioritized genes in biological context (Fig. 1; Online Methods). Specifically, biological information of each input gene is provided to gain insight into previously associated diseases as well as drug targets by mapping OMIM^14^ ID and DrugBank^15^ ID. Tissue specific expression patterns for each gene are visualized as an interactive heatmap, and provide information on whether a gene is expressed in a certain tissue. Overrepresentation in sets of differentially expressed genes (DEG; sets of genes which are more (or less) expressed in a specific tissue compared to other tissue types) for each of 53 tissue types (Supplementary Table 3) based on GTEx v6 RNA-seq data^16^ is also provided to identify tissue specificity of prioritized genes (Online Methods; Supplementary Table 3). Enrichment in biological pathways and functional categories is tested using the hypergeometric test against gene sets obtained from MsigDB^17^ and WikiPathways^18^.

To validate the utility of FUMA, we applied it to summary statistics of the most recent GWAS for Body Mass Index (BMI)^19^(see Online Methods). FUMA identified 95 lead SNPs (from 223 independent significant SNPs) across 77 genomic risk loci (Fig. 2 and Supplementary Table 4-6), in accordance with the original study. Functional mapping prioritized 151 unique genes; 23 genes with deleterious coding SNPs (positional mapping), 128 genes with eQTLs that potentially alter expression of these genes (eQTL mapping), and 16 genes that had both deleterious coding SNPs and eQTLs (Supplementary Table 7). The 151 genes include 55 genes that were also reported in the original study^19^ and 96 novel genes implicated by FUMA (Fig. 2). These novel candidates have shared biological functions with the 55 previously known candidate genes such as ‘metabolism of carbohydrate’, ‘metabolism of lipid and lipoprotein’, ‘immune system’ and ‘calcium signalling’ (Supplementary Table 8). In addition, the FUMA results showed that, although several genomic loci for BMI included multiple prioritized genes, a single gene was prioritized in 22 loci, suggesting that these 22 genes have a high probability of being the causal gene in that region. The 22 ‘highly likely causal genes’ include several well-known genes for BMI such as *NEGR1*, *TOMM40* and *TMEM18* (Supplementary Fig.2 and Supplementary Table 7). The strongest GWAS association signal for BMI was on 16q.12.2 where 3 genes were prioritized; *FTO*, *RBL2* and *IRX3* (Fig. 3). These three genes were only prioritized by eQTL mapping as the positional mapping showed no deleterious coding SNPs located in these genes. The original study^19^ only mentioned *FTO,* because the associated SNPs were located in this gene, however none of the associated SNPs have a potential direct affect such as coding SNPs on *FTO*. Two of the genes prioritized by FUMA (*RBL2* and *IRX3*) are physically located outside the genomic locus and are missed when using conventional approaches that prioritize genes located in the locus of interest based on LD around the top SNP. Although the *IRX3* gene was not reported in the original study^19^, recent functional work has indeed validated this as the causal gene whose expression is affected by SNPs in the 16q.12.2 locus^20^. To assess whether the prioritized genes converge on biological shared functions or pathways, FUMA tested for enrichment in GO terms, and canonical pathways. 15 significantly enriched GO terms were detected, including known and novel pathways, e.g. ‘Zinc ion homeostasis’ and ‘Glutathione related biological processes’ (Supplementary Table 10). Thus, using BMI summary statistics, FUMA confirmed known genes but also prioritized novel genes, including potential causal genes located *outside* the GWAS risk loci of BMI, which were missed in the original study.

**Figure 2.**
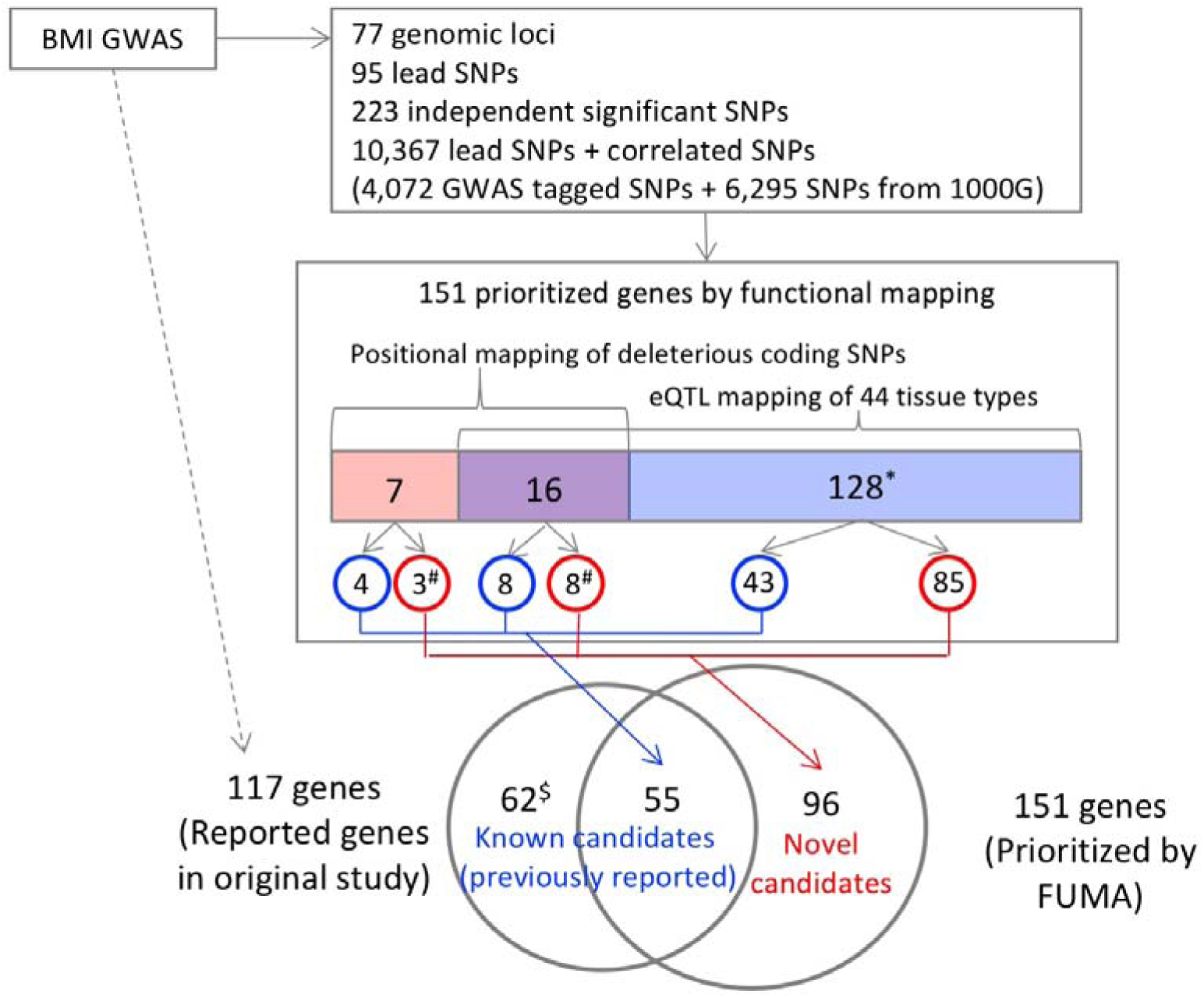
Overview of prioritized genes from BMI GWAS by FUMA. Starting from the BMI GWAS summary statistics, boxes represent results of the *SNP2GENE* process. The annotated SNPs include all independent lead SNPs and SNPs which are in LD with these lead SNPs. Prioritized genes are divided into three categories; genes that are implicated by deleterious coding SNPs (colored in pink), by eQTLs for these genes (colored in blue), or genes implicated by both strategies (colored in purple). The prioritized genes are further categorized into previously reported genes (blue circles) and novel genes (red circles) prioritized genes by FUMA. ^*^50 of the 128 genes are located outside of GWAS risk loci. ^#^These genes are located within the GWAS risk loci (since they have coding SNPs) but were not reported in the original study because of the following reasons: 1) FUMA considers all independent significant SNPs while only top SNPs were considered in the original study, or 2) FUMA incorporates non-GWAS tagged SNPs which are in LD of independent significant SNPs, or 3) reported genes do not necessary include all genes that are located within GWAS risk loci because the authors only choose to highlight a subset. ^$^These genes were not prioritized by functional mapping since they do not have either deleterious coding SNPs or eQTLs, although they are located within GWAS risk loci.

**Figure 3.**
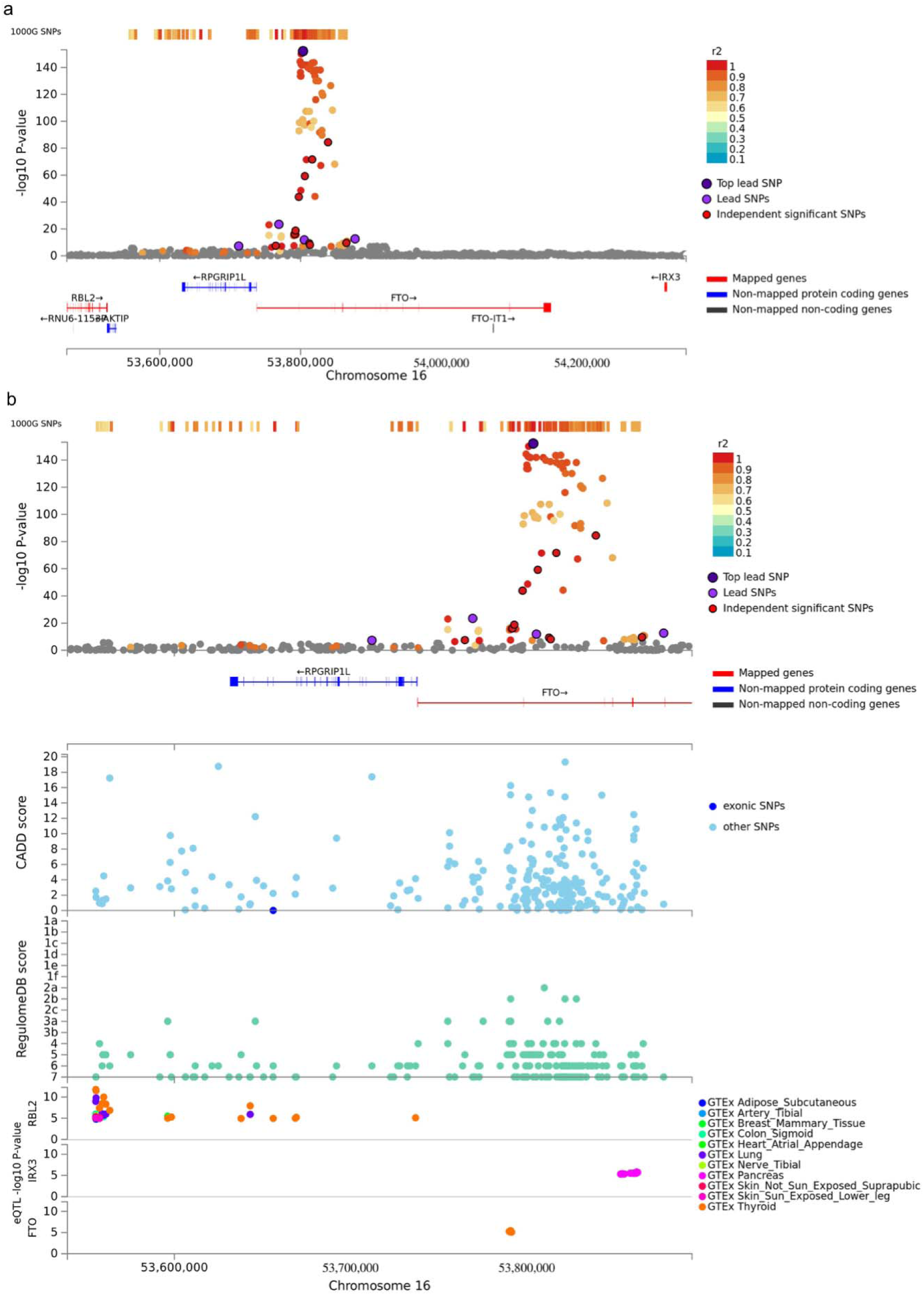
Regional plot of the locus 16q.12.2 of BMI GWAS and prioritized genes. (a) Extended region of the FTO locus, which includes prioritized genes *RBL2* and *IRX3*. Genes prioritized by FUMA are highlighted in red. (b) Zoomed in regional plot of FTO locus with, from the top, GWAS P-value (SNPs are colored based on r^2^), CADD score, RequlomeDB score and eQTL P-value. Non-GWAS-tagged SNPs are shown in the top of the plot as rectangles since they do not have a P-value from the GWAS, but they are in LD with the lead SNP. eQTLs are plotted per gene and colored based on tissue types. From these results, it can be seen e.g. that SNPs that were not originally included in the GWAS, but are known to be in LD with the lead SNP using the 1000 genomes reference panel, influence expression of *RBL2* in several different tissues. In addition, GWAS SNPs with a significant BMI association P-value and which are located in the FTO gene act as eQTL for expression of *IRX3* in the pancreas. The web-based version of this plot is interactive and allows zooming in or out as well as obtaining specific details about single SNPs.

To further illustrate its utility, we applied FUMA to the summary statistics of two other traits: Crohn’s disease^21^ (CD) and Schizophrenia^3^ (SCZ) (see Online Methods), where we obtained similar results: FUMA confirmed several genes that were reported in the original study, yet also prioritized genes that had not previously been reported (see Supplementary Results for details). For every prioritized gene, FUMA provides the reason for pinpointing this gene, such as for example when the expression of the prioritized gene is altered by a SNP that is in LD with or associated with the disease of interest. Interactive regional plots (Supplementary Fig. 5-7, 10-11) show which genes in a genomic risk locus are prioritized and which genes are not, and the annotated SNPs in the prioritized genes facilitate the generation of hypotheses for functional validation experiments. For example, if a gene is prioritized because of an associated loss-of-function SNP, follow-up validation experiments focusing on a knock-out of this gene may provide disease relevant functional information. On the other hand, if a gene is prioritized because a risk associated allele of a SNP increases expression of this gene in brain, then an overexpression experiment of this gene in neuronal cell cultures would be a more relevant experiment.

In summary, FUMA provides an easy-to-use tool to functionally annotate, visualize, and interpret results from genetic association studies and to quickly gain insight into the directional biological implications of significant genetic associations. FUMA combines information of state-of-the-art biological data sources in a single platform to facilitate the generation of hypotheses for functional follow-up analysis aimed at proving causal relations between genetic variants and diseases.

## ONLINE METHODS

### Data Sources and Pre-processes

Data repositories and tools used in FUMA are available in Supplementary Table 1. All genetic data sets used in this study are based on the hg19 human assembly and rsIDs were mapped to dbSNP build 146 if necessary. To compute minor allele frequencies and LD structure, we used the data from the 1000 Genomes Project^22^ phase3. Minor allele frequency and r^2^ of pairwise SNPs (up to 1Mb apart) were pre-computed using PLINK^23^ for each of available populations (AFR, AMR, EAS, EUR and SAS). Functional annotations of SNPs were obtained from the following three repositories; CADD^9^, RegulomeDB^10^ and core 15-state model of chromatin^
11–13^. Cis-eQTL information was obtained from the following 4 different data repositories; GTEx portal v6^16^, Blood eQTL browser^24^, BIOS QTL Browser^25^ and BRAINEAC^26^ and genes were mapped to ensemble gene ID if necessary. Genomic coordinate of GWAS catalog^1^ reported SNPs was lifted down using liftOver software from hg38 to hg19. Normalized gene expression data (RPKM, Read Per Kilo bae per Million) from GTEx portal v6^16^ for 53 tissue types were processed for different purposes. The details are described in ‘GTEx Gene Expression Data Set’ section. Curated pathways and gene sets from MsigDB v5.2^17^ and WikiPathways^18^ which are assigned entrez ID.

### Characterization of genomic risk loci based on association summary statistics (step 1 in SNP2GENE)

To define genomic loci of interest to the trait based on provided GWAS summary statistics, pre-calculated LD structure based on 1000G of the relevant reference population (EUR for BMI, CD and SCZ) is used. First of all, independent significant SNPs which have the genome-wide significant P-value (≤ 5e-8) and independent from each other at r^2^ 0.6. For each independent significant SNP, all known (i.e. regardless of being available in the GWAS input) SNPs that have r^2^ ≥ 0.6 with one of the independent significant SNPs are included for further annotation (candidate SNPs). These SNPs may thus include SNPs that were not available in the GWAS input, but are available in the 1000G reference panel and are in LD with an independent significant SNP. Candidate SNPs can be filtered based on a user defined minor allele frequency (MAF ≥ 0.01). Based on the identified independent significant SNPs, lead SNPs were defined if they are independent from each other at r^
2
^ 0.1. Additionally, if LD blocks of independent significant SNPs are closely located to each other (less than 250kb, distance if based on the most right and left SNPs from each LD block), they are merged into one genomic locus. Each genomic locus can thus contain multiple independent significant SNPs and lead SNPs.

Besides using FUMA to determine lead SNPs based on GWAS summary statistics, users can provide a list of pre-defined lead SNPs. In addition, users can provide a list of pre-defined genomic regions to limit all annotations carried out in FUMA to those regions.

### Annotation of candidate SNPs in genomic risk loci (step 2 in *SNP2GENE*)

Functional consequences of SNPs on genes are obtained by performing ANNOVAR^8^ (“gene based annotation”) using Ensembl genes (build 85). Note that SNPs can be annotated to more than one gene in case of intergenic SNPs which are annotated to the two closest up and down-stream genes. CADD, RegulomeDB score and 15-core chromatin state are annotated to all SNPs in 1000G phase 3 by matching chromosome, position, reference and alternative alleles. eQTLs are also extracted by matching chromosome, position and alleles for each user selected tissue types, wherein SNPs can have multiple eQTLs for distinct genes and tissue types. Information on previously known SNP-trait associations reported in the GWAS catalog is also retrieved for all SNPs of interest by matching chromosome and position.

### Gene Mapping (step 3 in *SNP2GENE*)

Gene annotation is based on Ensembl genes (build 85). To match external gene IDs, we mapped ENSG ID to entrez ID yielding 35,808 genes which consist of 19,436 protein-coding genes, 9,249 non-coding RNA and other 7,123 genes (e.g. pseudogenes, processed transcripts, immunoglobulin genes and T cell receptor genes).

Positional mapping is performed based on annotations obtained from ANNOVAR^8^ for which we provide two options; maximum distance from SNPs to genes and functional consequences of SNPs on gene. When the former option is defined, FUMA maps SNPs to genes based on ANNOVAR annotation and a user defined maximum distance is applied for intergenic SNPs. Note that ANNOVAR prioritize an annotation of SNPs which are located in a genomic region where multiple genes are overlapped. For these SNPs, they are mapped to the annotated gene by ANNOVAR. When the latter option is provided, FUMA maps only SNPs which have selected annotations annotated by ANNOVAR.

For eQTL mapping, all independent significant SNPs and SNPs in LD of them are mapped to eQTLs in user defined tissue types. By default, only significant SNP-gene pairs (FDR ≤ 0.05) are used. Optionally, eQTLs can be filtered based on a user defined P-value. eQTL mapping maps SNPs to genes up to 1Mb apart (cis-eQTLs).

Optional filtering of SNPs based on functional annotations obtained in step 2 of *SNP2GENE* (i.e. CADD score, RegulomeDB score, 15-core chromatin state) can be performed for positional and eQTL mappings separately.

### Functional Mapping: identification of potential causal genes from functional SNPs

We refer to “functional mapping” as the combination of positional mapping of deleterious coding SNPs, and tissue specific eQTL mapping. With functional mapping, we aim to further identify candidate causal genes based on biological function of SNPs. We include deleterious coding SNPs, either being exonic or splicing with CADD score ≥ 12.37 (defined by Kircher *et al.*
^
9
^), and eQTLs of defined tissue types (FDR ≤ 0.05).

### MAGMA: Gene Analysis and Gene set Analysis

In FUMA, input GWAS summary statistics is used to compute gene-based P-values (gene analysis) and gene set P-value (gene set analysis) by MAGMA^
27
^ to provide a genome-wide distribution of genetic associations. For gene analysis, the gene-based P-value was computed for protein-coding genes by mapping SNPs to genes if SNPs are located within the genes. For gene set analysis, the gene set P-value was computed using gene-based P-value for 4,728 curated gene sets (including canonical pathways) and 6,166 GO terms obtained from MsigDB v5.2. For both analyses, the default setting (SNP-wise model for gene analysis and competitive model for gene set analysis) were used, and the Bonferroni correction (gene) or False Discovery Rate (gene-set) was used to correct for multiple testing.

### GTEx Gene Expression Data Set

Normalized gene expressions (Reads Per Kilo base per Million, RPKM) of 53 tissue types were obtained from GTEx (Supplementary Table 3). A total of 56,320 genes was available in GTEx, which we filtered on an average RPKM per tissue greater or equal to 1 in at least one tissue type. This resulted in transcripts of 28,520 genes, of which 22,146 were mapped to entrez ID (see ‘Gene Mapping’ section for details). In the *GENE2FUNC*, the heatmap of prioritized genes displays two optional expression values; *i.* the average log2(RPKM+1) per tissue per gene, wherein RPKM was winsorized at 50, which allows comparison of expression level across genes and tissue types and *ii.* the average of the normalized expression (zero mean of log2(RPKM+1)) per tissue per gene which allows comparison of expression level across tissue types within a gene.

To obtain differentially expressed gene (DEG; genes which are significantly more or less expressed in a given tissue compared to others) sets for each of 53 tissue type, the normalized expression (zero mean of log2(RPKM+1)) was used. Two-sided Student’s t-tests were performed per gene per tissue against all other tissues. After the Bonferroni correction, genes with corrected p-value ≤ 0.05 and absolute log fold change ≥ 0.58 were defined as a DEG set in a given tissue, i.e. for these gene expression in the given tissue had the largest discrepancy with expression in all other tissues. In addition, we distinguished between genes that were up and down-regulated in a specific tissue compared to other tissues, by taking the sign of t-score into account. In *GENE2FUNC*, genes are tested against those DEG sets by hypergeometric tests to evaluate if the prioritized genes (or a list of genes of interest) are overrepresented in DEG sets in specific tissue types.

### Gene Set Enrichment Test

To test for overrepresentation of biological functions of prioritized genes, the prioritized genes (or a list of genes of interest) are tested against gene sets obtained from MsigDB (i.e. hallmark gene sets, positional gene sets, curated gene sets, motif gene sets, computational gene sets, GO gene sets, oncogenic signatures and immunologic signatures) and WikiPathways, using hypergeometric tests. The set of background genes (i.e. the genes against which the set of prioritized genes are tested against) is 19,264 protein-coding genes. Background genes can also be selected from gene types as described in ‘Gene Mapping’ section. Custom sets of background genes can also be provided by the users. Multiple testing correction (i.e. Benjamini-Hochberg by default) is performed per data source of tested gene sets (e.g. canonical pathways, GO biological processes, hallmark genes). FUMA reports gene sets with adjusted P-value ≤ 0.05 and the number of genes that overlap with the gene set > 1 by default.

### Validation with BMI GWAS

GWAS summary statistics for the BMI GWAS were obtained from http://portals.broadinstitute.org/collaboration/giant/index.php/GIANT_consortium_data_files and were used as input for FUMA. Parameters were set as described in the ‘Functional mapping’ section and we used eQTLs in 44 tissue types from GTEx. Indels were excluded. rsID was mapped to dbSNP build 146 and chromosome and positions were extracted based on human genome hg19 reference. Only protein-coding genes were used in gene mapping and enrichment of DEG in 53 tissue types, Canonical Pathways and GO terms were tested.

### Application to CD GWAS

GWAS summary statistics of CD was obtained from http://ftp.sanger.ac.uk/pub/consortia/ibdgenetics/. We set parameters as described in the ‘Functional Mapping’ section and we used eQTLs in 5 tissue types from GTEx which are relevant to CD, i.e. Small Intestine, Colon Sigmoid, Colon Transverse, Stomach and Whole Blood. The MHC region was excluded from the analysis. Since the input GWAS summary statistics only contained results from the discovery phase, we manually submitted the 71 reported lead SNPs to FUMA in addition to the independent lead SNPs that were identified as described above (Supplementary Table 11). Only protein-coding genes were used in mappings and enrichment of DEG in 53 tissue types, Canonical Pathways and GO terms were tested.

### Application to SCZ GWAS

GWAS summary statistics were obtained from http://www.med.unc.edu/pgc/results-and-downloads. Parameters were set as described in the ‘Functional mapping’ section, and eQTLs in 10 brain tissues from GTEx. The extended MHC region (25Mb – 34Mb), Chromosome X and indels were excluded from this analysis. The input GWAS summary statistics are based on the discovery phase and not all reported lead SNPs from the combined results of discovery and replication phasesreached genome-wide significance. To include all reported lead SNPs, 111 non-indel lead SNPs were provided to FUMA and additional independent lead SNPs were identified at P≤5e-8 (Supplementary Table 19). Only protein-coding genes were used in mappings and enrichment of DEG in 53 tissue types, Canonical Pathways and GO terms were tested.

## ACKNOWLEDGEMENTS

This work was funded by The Netherlands Organization for Scientific Research (NWO VICI 453-14-005) and Ingrosyl. We thank the GIANT consortium, WTCCC and PGC for providing GWAS summary statistics and GTEx Portal for RNA-seq and eQTL data.

## AUTHOR CONTRIBUTIONS

D. P. conceived the study. K. W. and A.v. B. developed the web application. K. W. performed analyses and drafted the manuscript. K. W., E. T., and D. P. participated in the discussions, interpretation of the results, and editing of the manuscript. All authors provided relevant input at different stages of the project and approved the final manuscript.

## COMPETING FINANCIAL INTERESTS

The authors disclose no potential conflicts of interest.

